# *In silico* modelling of CD8 T cell immune response links genetic regulation to population dynamics

**DOI:** 10.1101/2024.03.01.582928

**Authors:** Thi Nhu Thao Nguyen, Madge Martin, Christophe Arpin, Samuel Bernard, Olivier Gandrillon, Fabien Crauste

**Affiliations:** Univ Lyon, ENS de Lyon, Univ Claude Bernard, CNRS UMR 5239, INSERM U1210, Laboratory of Biology and Modelling of the Cell, 46 alĺee d’Italie Site Jacques Monod, F-69007, Lyon, France; Inria, Villeurbanne, 69693, France; CNRS, Univ Paris Est Creteil, Univ Gustave Eiffel, UMR 8208, MSME, F-94010 Cŕeteil, France; Centre International de Recherche en Infectiologie, Université de Lyon, ENS de Lyon, Université Claude Bernard, CNRS UMR 5308, INSERM U1111, Lyon, France; Univ Lyon, Université Lyon 1, CNRS UMR5208, Institut Camille Jordan, 43 Blvd du 11 Novembre 1918, Villeurbanne-Cedex, F-69622, France; Université Paris Cité, CNRS, MAP5, F-75006 Paris, France

**Keywords:** ***Keywords—*** Gene Regulatory Networks, cell population dynamics, CD8 T cells, stochastic gene expression, multiscale modeling

## Abstract

The CD8 T cell immune response operates at multiple temporal and spatial scales, including all the early complex biochemical and biomechanical processes, up to long term cell population behavior.

In order to model this response, we devised a multiscale agent-based approach using Simuscale software. Within each agent (cell) of our model, we introduced a gene regulatory network (GRN) based upon a piecewise deterministic Markov process (PDMP) formalism. Cell fate – differentiation, proliferation, death – was coupled to the state of the GRN through rule-based mechanisms. Cells interact in a 3D computational domain and signal to each other via cell-cell contacts, influencing the GRN behavior.

Results show the ability of the model to correctly capture both population behaviour and molecular time-dependent evolution. We examined the impact of several parameters on molecular and population dynamics, and demonstrated the add-on value of using a multiscale approach by showing that a higher degradation rate for the protein controlling cell death induces a later peak in the response.

## 1 Introduction

CD8 T cells are important for immune responses against viruses and intracellular bacteria, as well as for tumor surveillance. A naive CD8 T cell gets activated when it recognizes an antigen presenting cell (APC), through the formation of an immunological synapse [16]. Activated CD8 T cells first give rise to proliferating memory precursor (MP) cells [54]. Such MP cells could represent bipotential cells that face a choice between two fates: the terminally differentiated effector fate that is associated with the repression of their self-renewing capacity and the activation of their effector function, and the memory precursor fate that maintains their self-renewing capacity. Effector T cells massively proliferate while acquiring their effector potential, which allows them to kill infected or antigen-bearing malignant cells, before dying during the contraction phase. Meanwhile, part of MP cells differentiate into memory cells, providing long-term protection against reinfection [29] The capacity to generate long-lived memory cells during a primary immune response forms the basis for vaccination. At the end of the response, the number of memory cells remains stable, since memory cells hardly ever proliferate or die [13, 39].

This entire process functions across various temporal and spatial scales, encompassing intricate early biochemical and biomechanical processes, extending to the long-term behavior of cell populations. Building a comprehensive computational model, where all relevant scales and their interactions would be represented, would hold the potential to introduce new and more robust principles for designing vaccines aimed at swiftly adapting pathogens. This perspective motivated the recent development of advanced mathematical and computational models to depict these multiscale phenomena, moving us towards a more comprehensive understanding of the CD8+ T cell response [5].

There is a rich ecosystem of software developed to describe cell dynamics systems at different scales.

Software can be distributed into three main classes:

1. Description of cell population dynamics, such as Compucell3D [52], Physicell [19] or CellSys [23].
2. Description of both cellular and intracellular scale dynamics, such as Virtual Cell [12], COPASI [24] or Smoldyn [3].
3. Description of coupling at least two different scales, like Vivarium [1], ENISI-MSM [59], EpiLog [56] coupled with COPASI, MSM [53] or Tissue Forge [48]. One should also cite PhysiBoSS [36] which results from the coupling of Physicell (processing up to 10^6^ cells, but needs to run for several days) with MaBoSS, a tool based on Boolean modeling [10].

These tools are limited in some aspects for example in the number of cells they can simulate, the explicit description of a molecular level or their ability to deal with different cell types. Their computation time is also often extremely slow, limiting the relevance of such computational models for performing parameter estimation and model fitting, at least when dealing with highly proliferating cells. Indeed, when the number of cells is only a few thousands, the calculation process becomes cumbersome and time-consuming.

The development of a multiscale model of the immune response requires the ability to simulate the molecular state of an expanding T cell population over several scales at the single-cell level, using a realistic GRN that can reproduce stochastic gene expression behavior.

In [18, 44], CompuCell3D was coupled with a molecular network described by an ODE system. Although these latest studies have qualitatively captured expected cellular and molecular behaviors, enabling cellular decision-making, the molecular network was modeled as a fully deterministic system using ODEs, whereas it is now accepted that gene expression at the single cell level is a stochastic process [7, 11, 17, 45, 46, 49, 51].

We therefore describe here the use of Simuscale [8], which enabled us to simulate the dynamics of interactions from the molecular to the cell-population scale, in conjunction with the use of a biologically realistic mechanistic GRN based on a stochastic 2-state model for gene expression [22]. Simuscale is particularly relevant to model and simulate differentiating cell populations, whose dynamics are not solely dependent on biophysical rules. Thanks to appropriate parameter calibration, this study demonstrates the ability to capture the expected time-dependent evolution of CD8 T cell population dynamics. In addition, issues related to simulation time, variability, the dependence on the initial condition of the number of cells in the population and differentiation states, were also addressed in this study. Finally, we demonstrate the benefit of multiscale coupling by assessing the population behavior (time of the peak of the response) as a function of a molecular parameter (protein half-life).

## 2 Material, Methods and Models

### 2.1 Simuscale

Simuscale [8] is a multiscale individual-based modeling platform for performing numerical simulations of heterogeneous populations of individual cells. Cells are assumed to evolve in time and interact physically and biochemically with each others. Models are described at two levels: cellular and population level. The cellular level describes the dynamics of single cells, as defined by the user/modeler. Cells have an internal state that includes default properties such as cell size and position, and may also include cell-specific states (*e.g.*, gene or protein expression). The population level describes the mechanical constraints and biochemical interactions between cells. Cells evolve in a bounded 3D domain, and can divide or die. Cells are represented spatially as visco-elastic spheres with a rigid core.

Simuscale implements the physical simulator that manages the simulations at the population level. Details of cellular dynamics to each cell are to be defined by the user. This makes Simuscale modular, as it can accommodate any number of cell models within the same simulation, including models with different modelling formalisms, such as ordinary and stochastic differential equations and up to Piecewise Deterministic Markov Process (PDMPs, [22]). Biochemical interactions occur between cells that are in contact with each other, through intercellular signals. Intercellular signals can be known to all or to a subset of the cells only.

Simuscale expects an input file describing the initial cell population and numerical options. It runs a simulation over a specified time interval, updating the cell population at given time steps, and it generates an output file containing the state of each cell at each time step, and the tree of cell divisions and deaths. All details regarding Simuscale can be found in Bernard et al. [8]

### 2.2 Gene Regulatory Network

The first step in building a multiscale model of the CD8 T cell immune response is to build a GRN whose dynamics will drive each CD8 T cell fate (proliferation, death, differentiation).

We first build a simplified GRN, based on essential principles that allow each cell to be able to proliferate, die, and differentiate in each relevant cell type (mainly, effector or memory CD8 T cell), see Section 3.1. This GRN is made of 9 genes, which have no equivalent in a real-biological setting (yet analogies are drawn in Section 3.2), each gene dynamics being driven by the gene model introduced in Section 2.3.

In order to obtain realistic dynamics, we also consider an augmented GRN based on the 9-genes GRN, to which three so-called ‘decorating genes’ are added to each main gene, resulting in a 36-genes GRN with the same properties. A “decorating gene” is activated by its main gene but does not act on any downstream gene and therefore has no influence on the GRN dynamics.

The GRNs are dynamical mathematical models, based on the coupling of deterministic and probabilistic formalisms, resulting in stochastic dynamics (see Section 2.3). In particular, genes of the GRN can act on other genes of the GRN, either by activating or inhibiting their expression, resulting in highly nonlinear dynamics.

It is important to note that each CD8 T cell will be embedded with the same GRN. Nonetheless, depending on their previous experiences and interactions with other cells, CD8 T cell molecular, intra- cellular states will differ from cell to cell and will be specific to each cell, thanks to individual values of each gene expression in a given cell.

### 2.3 Gene model

Given the stochastic nature of gene expression at the single cell level [17, 32], we chose to model the expression of each gene in the GRN (see Section 2.2) as a stochastic two-state model [42]. This stochastic process consists of three components for each gene *i*, *i ∈ {*1*, …, n}*: the promoter state *E_i_*, the mRNA level *M_i_*and the protein level *P_i_*.

The promoter can be in two states *E_i_* = 0 or 1 (inactive or active). It opens with a rate *k*_on_ and closes with a rate *k*_off_ . When the promoter is in the open state, mRNA gets synthetized at a rate *s*_0_ and degraded at a rate *d*_0_. From mRNA, proteins get synthetized at a rate *s*_1_ and degraded at a rate *d*_1_.

Such a model can be implemented in a variety of formalisms [21]. In the present work we consider the so-called bursty regime of the two-state model. It corresponds to the experimentally observed situation where active periods are short and characterised by a high transcription rate, thereby generating bursts of mRNA [51]. In such a regime, the promoter state no longer needs to be explicitly described since active periods can be considered as infinitely short: random jumps will instantaneously increase the amount of mRNA levels *M_i_*.

The construction of a gene trajectory is as follows: starting from state (*M_i_*(*t*)*, P_i_*(*t*)), the dynamics of mRNA and protein levels are given by

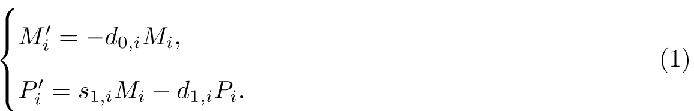

until a burst occurs for an isolated gene *i*, at rate *k*_on_*_,i_*. Following the burst, the quantity *M_i_* jumps by a random height according to an exponential law with rate *k*_off_*_,i_/s*_0_*_,i_* (see [21] or [57] for details of the model). This stochastic model constructs a piecewise-deterministic Markov process (PDMP).

To describe regulatory mechanisms within the GRN, we introduce the dependence of the burst frequency upon the proteomic field and the external signalling activity, i.e. a burst occurs for gene *i*, at a rate *k^θ^* (*P, S*), where *P* = (*P*_1_*, . . . , P_n_*) is the protein level vector of all genes in the network and *S* = (*S*_1_*, . . . , S_m_*) is the signalling state vector of the network. For every *i ∈* (1*, . . . , m*), *S_i_*= 1 or 0 (active or inactive signalling). More precisely, in [21, 57], the burst rate of each gene *i* is calculated using

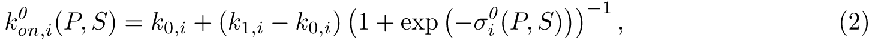

where *k*_0*,i*_ and *k*_1*,i*_ correspond respectively to the minimum and maximum burst frequencies of gene *i*, and

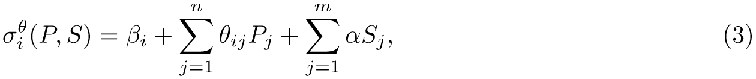

where *β_i_* is the basal activity of gene *i* [21], *α* is a signalling parameter, and *θ* = *{θ_ij_}_i,j∈{_*_1_*_,…,n}_* is the gene-gene interaction matrix, representing the considered GRN (see Supplementary Figure 1). Parameters *θ_ij_*can be positive or negative corresponding either to the activation or the inhibition behaviour.

### 2.4 Cell types

When a simulation starts, there are two cell types in the population, APC and naive CD8 T cell. APC do not possess an internal state, whereas we recall that all CD8 T cells share the same GRN structure and CD8 cell types are defined as a function of the dynamical state of their GRN. A naive cell becomes activated when it contacts an APC for a sufficiently long period of time. This “activated” state will be transmitted throughout the descent of an activated cell. Activated CD8 T cells will then acquire specific cell type identity as a function of their position in the gene expression space. They will be considered bipotent if their protein levels in both genes 7 and 8 (see Section 3.1) are below certain threshold values *P ^∗^* and *P ^∗^*, respectively. When *P*_7_ *≥ P ^∗^* while *P*_8_ *< P ^∗^*, they will be considered effector cells, and when *P*_8_ *≥ P ^∗^*, they will be considered memory cells. Table 1 summarizes all CD8 T cell differentiation states considered in this work.

**Table 1:** Differentiation states of CD8 T cells.

| Cell type | Definition |
| --- | --- |
| Naive | No contact with an APC (inactivated) |
| Bipotent | Activated, protein levels $P_7 < P_7^*$, $P_8 < P_8^*$ |
| Effector | Activated, protein level $P_7 \geq P_7^*$, $P_8 < P_8^*$ |
| Memory | Activated, protein level $P_8 \geq P_8^*$ |

### 2.5 Signalling

We consider two kinds of signals, namely APC signalling (*via* T cell-APC contact) and TCC signalling (effector-T cell contact), inducing either activation or apoptosis in a CD8 T cell after its encounter with an APC or an effector cell, respectively (see Figure 1).

**Figure 1:**
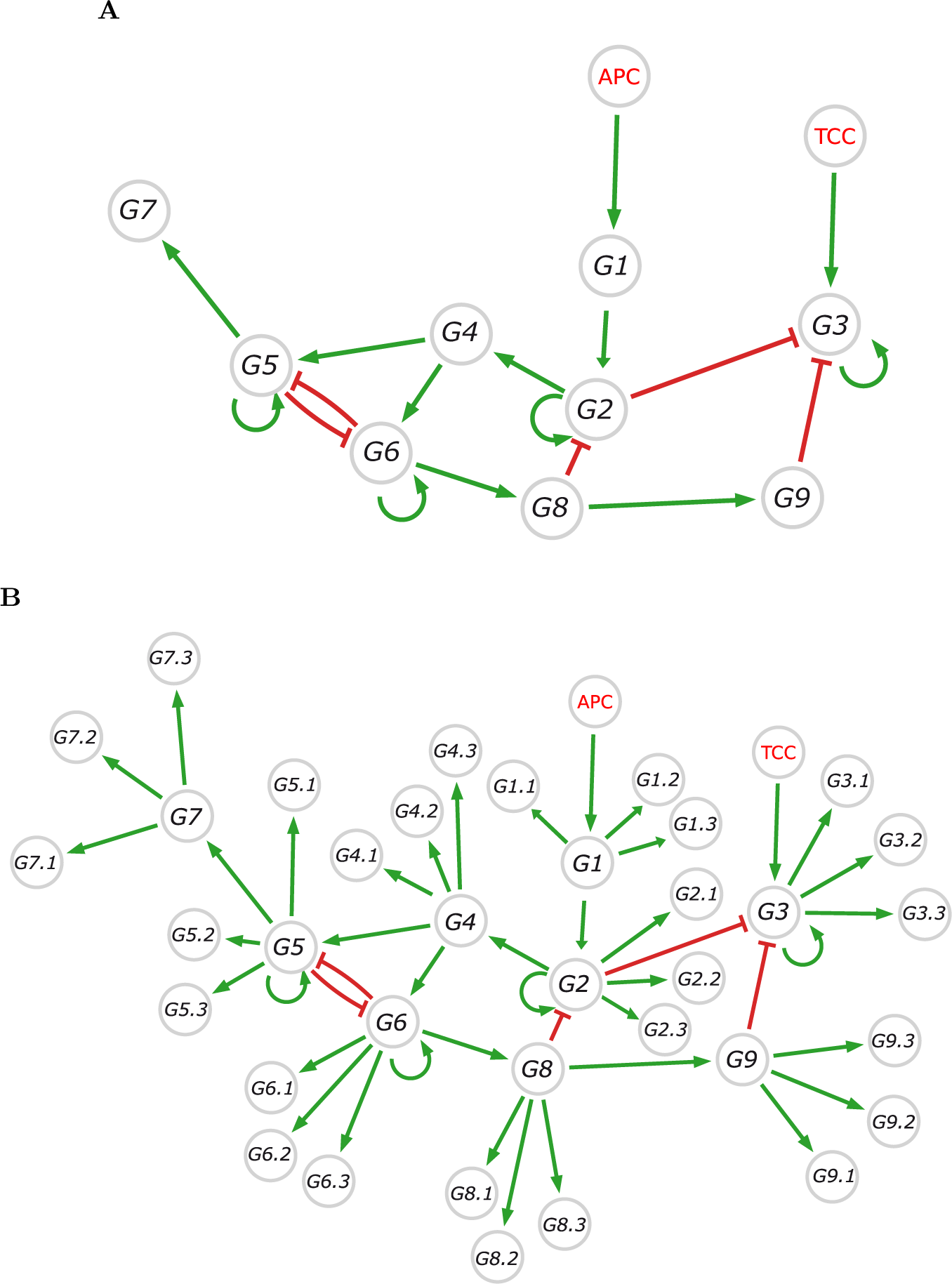
Schematic representation of the GRN. (A) The core circuitry of the principle-based GRN using 9 genes. The notation *Gi* corresponds to the gene number *i*. APC signalling and TCC signalling are perceived when CD8 T cells are in contact with APCs or effector cells, respectively. The green (resp. red) arrows represent activation (resp. inhibition). (B) The augmented GRN, made of 36 genes, incorporates the core principle-based GRN together with “decorating” genes. The gene *Gi.j*, *j ∈ {*1, 2, 3*}*, is the *j*-th gene activated by gene *i*.

A CD8 T cell perceives a signal from another cell (either an APC or an effector T cell) if

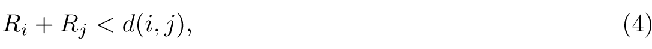

where *R_i_* and *R_j_* are the external radii of both cells and *d*(*i, j*) is the Euclidean distance between the centers of the two cells. Condition (4) means that a CD8 T cell comes into contact with another cell if their spheres intersect. Once a contact is detected, the phenotype of the contacting cell is determined and the corresponding signalling (APC signalling or TCC signalling) is applied.

When a naive CD8 T cell encounters an APC (contacts on its surface), it receives an APC signalling, which impacts the GRN behaviour (see below). It has been shown that when a CD8 T cell encounters an APC, it adheres tightly to the APC for up to 20 hours in *vitro* before starting to proliferate [28]. Therefore, in the model, until the first proliferation occurs an activated T cell remains attached to an APC, through adjusting the coefficient *σ_T_* (Table 5). More precisely, when a naive CD8 T cell encounters an APC, it tends to stay with this APC for a while thanks to a modification of its velocity coefficient *σ_T_* which takes a very low value. Then the APC signals to the T cell (APC signalling equals 1). If the activated CD8 T cell has not yet divided for the first time and the cumulative contact time with the APC is greater than 10 hours (i.e. it only takes 5 hours if the CD8 T cell encounters 2 APCs at the exact same time), gene number 1 is activated by the APC signaling (the level of expression of this gene determines the first proliferation, see Section 3.1), with a signalling parameter *α* = 20 (no unit) in order to strongly initiate gene expression. The effect of this value on model’s outputs has been numerically investigated and shows very limited impact (Supplementary Figure 2). Noticeably, a CD8 T cell may leave the APC before its protein 1 level reaches the value required for first division (this is rare, but possible due to the probabilistic nature of the model). In this case, due to its fast mobility (*σ_T_*goes back to its initial value upon contact breaking), the CD8 T cell will easily contact a new APC, its *σ_T_* value will decrease again and its protein level will increase again rapidly.

In all cases, immediately after its first division, the CD8 T cell leaves the APC and moves randomly. Further contacts with APC for this cell may occur, yet subsequent divisions will no longer depend on the APC signalling. In particular, this will be the case for the daughter cells of activated cells, as we assume that both daughter cells remember their mother’s first division, so they will not need to be activated by the APC signalling in order to proliferate [55].

Concerning apoptosis, we assume effector CD8 T cells’ targets include all CD8 T cells, so fratricide killing may occur [35, 50]. In this case, an effector cell sends an apoptotic signal called TCC signalling to another CD8 T cell it contacts. This leads to an increase in gene 3 bursting frequency (see Section 3.1), and protein 3 level increases in the receiver cell, eventually resulting in its death. Only an effector cell can emit TCC signalling. When a CD8 T cell receives a signal emitted by an effector cell, its TCC signalling value equals 1 and its signalling parameter *α* equals 20 (no unit).

### 2.6 Initial conditions

#### Initial molecular content

In order to obtain realistic (i.e. non-zero) values for the initial molecular content, we initialize each simulation with the values observed after 24 hours of simulating T cells in the absence of APCs. Specifically, we run the simulation with 980 CD8 T cells in the absence of APCs, so that they start to synthesize mRNAs and proteins without being activated. We then use simulated quantities of mRNA and proteins as initial data for the naive cells of the next simulation, where they are seeded in presence of APCs at time *t* = 0. Noticeably, APCs do not have a dynamical internal state, only CD8 T cells do.

#### Initial positions in the computational domain

Initial positions of APCs and CD8 T cells are uniformly randomized within the bounded computational domain (see Supplementary Figure 3).

### 2.7 Cell motion and fate

#### Cell movement

T cells move randomly in the computational domain. The random movement of T cells was modelled through the use of a Gaussian distribution. More precisely, at each time step *dt*, new 3D coordinates of CD8 T cells are updated by an amount identically distributed in each direction, as follows:

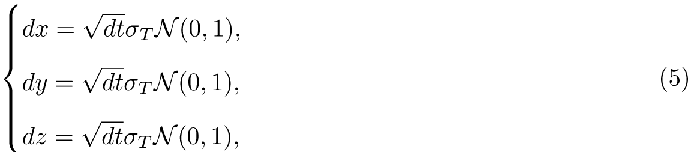

where *σ_T_*characterizes the velocity of the CD8 T cell when in contact or non-contact with an effector cell or an APC (see Table 5).

#### Cell fate

Regarding the proliferation process, if a CD8 T cell reaches its maximum volume (through the linear growth function of the current volume) and satisfies a condition on protein levels (*P*_1_ *≥ P ^∗^* for the first division, *P*_2_ *≥ P ^∗^* for subsequent divisions, see Section 3.1), then it divides into 2 daughter cells of equal volume. At division, the molecular content *C* of the mother cell is normally distributed between both daughter cells, and equals (1 *±* 0.2*N* (0,0.2))*C*/2 when *C* is the protein concentration and (1 *±* 0.1*N* (0, 0.2))*C/*2 when *C* is the mRNA concentration.

Concerning apoptosis, if a CD8 T cell reaches the condition of death (protein level of gene 3 is above a threshold value *P ^∗^*), then the cell disappears from the population immediately. We also set up an APC death mechanism to ensure that they will start dying after day 12 (i.e. after the expansion phase) and will disappear completely from the computational domain by day 20. This is a way to mimic the displacement of CD8 T cells out of the APC-containing organ [41], or the eradication of the antigen.

Cell differentiation is discussed in 2.4, see Table 1.

### 2.8 Simulation Parameters

All parameters used throughout this study are introduced in this section. Some parameters are needed for the functioning of the core code of Simuscale , other parameters are specific to the CD8 T cell simulation.

For running Simuscale simulations, we use an environment simulation domain inside a cube measuring 40x40x40 (same unit as the initialized radius, see [8]). This is what will be called ‘computational domain’ in this paper. Assuming cells are around 10 *µ*m in diameter, a space unit (SU) would correspond to around 7 *µ*m, for a domain side approximately 300 *µ*m in length. Each cell is a sphere with a maximum volume of 2 SU^3^ and a minimum volume of 1 SU^3^. There is a linear growth function, where the initial volume of each cell is randomly chosen in the range [1, 2], and the growth factor is fixed at 2*^dt/^*^10^, i.e. it takes 10 hours for a cell with a minimum volume to reach its maximum volume.

The time step of all simulations is set to *dt* = 0.1 hour, and the final simulation time is 960 hours, i.e. 40 days.

For T cell specific simulations, various parameters associated with cell fate, motion, and contacts must be defined. We will first focus on parameters related to gene dynamics.

The *β_i_* parameter represents the basal activity of gene *i*, in a logarithmic scale, in the absence of any stimulation (see equation (3)). Its value was arbitrarily fixed to *β_i_* = *−*5 for all genes *i* and allowed to reproduce consistent behaviors of gene dynamics. Values of the parameters of the gene-gene interaction matrix are shown in Table 2.

**Table 2:** Parameter values of the gene-gene interaction matrix. Parameters *θ_i,j_* give the value of the action of gene *i* onto gene *j*, while parameters *θ_i,i.j_* represent the interaction between gene *i* and its three decorating genes *i.j* (*j ∈ {*1, 2, 3*}*).

| Parameter | $\theta_{1,2}$ | $\theta_{2,2}$ | $\theta_{2,3}$ | $\theta_{2,4}$ | $\theta_{3,3}$ | $\theta_{4,5}$ | $\theta_{4,6}$ | $\theta_{5,5}$ | $\theta_{5,6}$ | $\theta_{5,7}$ |
| --- | --- | --- | --- | --- | --- | --- | --- | --- | --- | --- |
| Value | 40 | 16 | -10 | 16 | 60 | 16 | 16 | 90 | -150 | 25 |
| Parameter | $\theta_{6,5}$ | $\theta_{6,6}$ | $\theta_{6,8}$ | $\theta_{8,2}$ | $\theta_{8,9}$ | $\theta_{9,3}$ | $\theta_{i,i,j}$ | All other $\theta_{i,j}$ | | |
| Value | -150 | 90 | 15 | -800 | 530 | -3000 | 15 | 0 |  |  |

Values of protein degradation rates for all genes are given in Table 3. Since *d*_1_ *≪ d*_0_, we kept the degradation rate of mRNA equal to 1 for all genes, so *d*_0_*_,i_* = 1 for *i ∈ {*1*, . . . , n}*, and varied only the protein degradation rates. Those are chosen to progressively decrease along the main GRN, thus creating a delay in expressions and a wave-like behavior of signal propagation in the GRN [9]. Burst frequency of gene *i* is defined by *k*_off_*_,i_/s*_0_*_,i_*, whose value has been chosen to correspond to an exponential distribution of bursts with mean *µ* = 50. The effect of this value on model’s outputs has been numerically investigated and shows limited impact (Supplementary Figure 2).

**Table 3:** Gene kinetic parameters. Values of: protein degradation rates *d*_1_*_,i_* of gene *i* ; protein degradation rates *d*_1_*_,i.j_* of decorating genes *i.j*, *j ∈ {*1, 2, 3*}*, of gene *i* ; burst frequency *k*_off_*_,i_/s*_0_*_,i_* of gene *i* ; synthesis rates *s*_1_*_,i_* of gene *i* proteins (translation rates).

|  |  |  |  |  |  |  |
| --- | --- | --- | --- | --- | --- | --- |
| Protein degradation rate | $d_{1,1}$ | $d_{1,2}$ | $d_{1,3}$ | $d_{1,4}$ | $d_{1,5}$ | $d_{1,6}$ |
| Value | 0.03 | 0.021 | 0.021 | 0.02 | 0.019 | 0.009 |
| Protein degradation rate | $d_{1,7}$ | $d_{1,8}$ | $d_{1,9}$ | $d_{1,i,j}$ | | |
| Value | 0.02 | 0.00014 | 0.00013 | 0.03 |  |  |
| Synthesis rate | $k_{\text{off},i}/s_{0,i}$ | $s_{1,i}$ | | | | |
| Value | 0.02 | $0.01d_{1,x}$ | | | | |

Threshold values for proteins defining the cell types are given in Table 4.

**Table 4:**
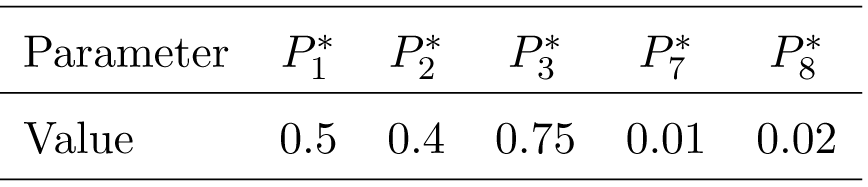
Protein threshold values. *P ^∗^* is the threshold value for the protein of gene *i*, used to define cell fate (proliferation, apoptosis) and differentiation states, see Table 1.

**Table 5:** CD8 T cell velocities.

| $\sigma_T$ | Contact behaviour with respect to cell velocity |
| --- | --- |
| 5 | no contact with others/after first division upon encountering APC |
| 0.02 | contact with an APC |
| 0.8 | contact with an effector cell |

Finally, Table 5 presents the velocity coefficients of CD8 T cells in different cases. The default behavior of CD8 T cells is to move randomly and fast. When in contact with another T cell, they reduce their speed, and almost stick to an APC when they first encounter the APC, until their first proliferation occurs. Consequently, three different velocity parameters are used depending on the situation.

### 2.9 Model’s outputs and simulations

All simulations are performed by running Simuscale code on a computation platform of the “P^ole Sci- entifique de Modélisation Numérique” (PSMN) of ENS Lyon. During each simulation, all cell population information is recorded, including the total number of cells in the population, cell identity, coordinates, volume, signalling, cell type and molecular content. This information is then post-processed on personal computers, using Python and R tools. All figures related to model’s outputs presented in this paper have been generated from the outputs of the simulations and plotted with Python and R.

Typical simulation times range from a few minutes (35 CD8 T cells and 25 APCs initially) to a few hours (1960 CD8 T cells and 1400 APCs initially ). It took only 2h to simulate the simulation associated to the largest initial cell counts in this study (average value over 10 simulations), executed on PSMN’s partition Cascade with 96 cores and 384 GB RAM (for more details, see https://www.ens-lyon. fr/PSMN/Documentation/clusters_usage/partitions_overview.html). All codes are available at https://gitlab.inria.fr/thinnguy/Simuscale_Lymphocytes

It is important to note that among the functionalities of Simuscale there is the possibility to color-code cells according to the quantitative value of any cellular variable. Coupled with a video display of he output, this is an invaluable feature of the model to:

1. Verify the correct functioning of the model. For example do divisions occur as planned when a certain value of *P*_2_ is reached, as expected?
2. For many biologists ”seeing is believing”. Videos are therefore the perfect medium for interactions between modellers and biologists.
3. In the present work there is no resulting 3D structure since cell movement tends to homogenize spatial cellular localisation, but for modeling spatially constrained structures, this output would be critical.

As an illustrative example, a movie of the CD8 T cell population dynamics is available at: https://gitlab.inria.fr/thinnguy/Simuscale_Lymphocytes/ (Pop T cell APC.mov). All relevant cellular variables are examined during the course of the response.

### 2.10 Sensitivity analysis

We perform a sensitivity analysis in Section 3.6. We consider two types of influence: either a sensitivity to initial conditions or to molecular parameters. In each case, we fix parameter values, perform 10 simulations, calculate the mean and standard deviations of the outputs, and compute variables of interest.

Variables of interest are: the number of CD8 T cells at the peak of the response; the ratio between the average number of CD8 T cells at the peak of the response and the initial number of CD8 T cells; the time of the response peak; the average number of memory CD8 T cells at the end of the simulation (*t* = 40 days); the ratio between the average number of memory CD8 T cells at the end of the simulation and the average number of CD8 T cells at the peak of the response.

The influence of initial conditions is assessed by either using different CD8 T cell and APC initial cell counts, but preserving the ratio T cell counts/APC counts (ratio equals to 1.4), or by fixing the initial number of CD8 T cells (two values were used, 210 and 980 cells) and varying the initial number of APCs (with T cell/APC ratios ranging in [0.3; 8.4]).

The influence of molecular content is assessed by computing variables of interest for various values of some molecular parameters, including: the degradation rates of proteins 2, 3, and 6 that determine proliferation, apoptosis and memory T cell differentiation, respectively. These parameters have been selected based on the *a priori* essential roles played by genes 2, 3 and 6 in controling CD8 T cell fate.

## 3 Results

### 3.1 A model involving a gene regulatory network constructed by 9 genes

For the intracellular scale, we first build a GRN composed of 9 genes (see Figure 1A), which triggers the proliferation, apoptosis, and differentiation of a single cell. All genes are hereafter denoted *Gi*, *i* = 1*, . . . ,* 9. From now on, we refer to this GRN as a ‘principle-based GRN’, since it is built upon few principles required to reproduce the main features of a CD8 T cell immune response. The logic behind this GRN is as follows.

First, APC signalling duration (cumulative contact time with APC is greater than 10 hours [28]) activates *G*1, which leads to the first proliferation of CD8 T cells when the protein concentration of *G*1 reaches a threshold value *P ^∗^* and the cell has doubled its volume (volume is equal to 2). After division, the activated CD8 T cell leaves the APC due to an increased velocity coefficient and the activation stops. As long as all APCs have not been eliminated, activated CD8 T cells can encounter APC again but they do not really attach to APC due to their high velocity.

In the meantime, *G*1 activates *G*2, which also activates itself in order to maintain a strong proliferation phase at the beginning of the immune response. The second and subsequent proliferation events are based on *G*2 dynamics: when the *G*_2_ protein level exceeds a threshold value *P ^∗^* then cells divide. Also, *G*2 inhibits *G*3, which induces CD8 T cell apoptosis above a threshold value *P ^∗^*. Gene *G*_3_ is activated through TCC signalling and *G*3 also activates itself.

Furthermore, *G*2 activates *G*4, which simultaneously activates two genes, *G*5 and *G*6. There is a toggle switch between these two genes which then creates the differentiation states. *G*5 activates *G*7, which will be the marker for the effector phenotype (when the protein of this gene exceeds a value *P ^∗^*). In addition, *G*6 activates *G*8, which induces the memory phenotype (when its protein concentration reaches a threshold value *P ^∗^*). Gene *G*8 inhibits *G*2 to mimic cell cycle exit of memory cells. Finally, *G*8 activates *G*9, which inhibits *G*3, leading to memory cell long-term survival. All threshold values are shown in Table 4.

For application purposes that require many more genes in the network (see Section 3.5), we also added so-called “decorating genes” to the principle-based GRN, resulting in a second GRN model (hereafter referred to as the ‘augmented model’). More precisely, each gene, from *G*1 to *G*9 of the principle-based GRN, simultaneously activates *n* redundant “decorating genes” that have no impact on the GRN dynamics. In the next sections, we illustrate the results with simulations of the augmented GRN, including three (*n* = 3) “decorating genes” for each main gene (see Figure 1B).

### 3.2 Biological rationale for our GRN

It is important to note that the principle-based GRN was built in order to satisfy minimal requirements imposed by a realistic description of the CD8 T cell immune response: it required genes able to induce proliferation, death and differentiation of CD8 T cells, therefore there is no reason any gene in this GRN has an equivalent specific gene in a real biological setting. Nevertheless, one could draw the following analogies between the principle-based GRN introduced previously and known genes.

1. *G*1 is activated by APC signalling and leads to the first cell division of activated cells. This has been shown to be a function carried by the *mTORC1* gene [37].
2. *G*2 drives the second and subsequent divisions. This role can be carried by the *IL-2R* gene [30], which also amplifies itself as IL-2 signalling promotes further IL-2R expression [15].
3. *G*3 leads to programmed cell death (apoptosis), which is known to be the activity of *Fas* [33]. In such a case, the TCC signalling could be seen as FasL signalling [40].
4. *G*5 and *G*6 are involved in a toggle switch, leading to the differentiation states. Two members of the T-box transcription factors have been demonstrated to play this critical role: *T-bet* (*G*5) [27] and *Eomesodermin* (*G*6) [26], which promote effector and memory differentiation, respectively. Similar tandems could be *Id2/Id3* and *Blimp1/Bcl-6* [29].
5. *G*7 is a marker for the effector phenotype. Many genes could fall in this category, but since in this model effector cells have the capacity to kill through the TCC signalling, *FasL* is one obvious candidate for this function [2].
6. *G*8 is a marker for the memory phenotype, limits growth and prevents death by apoptosis, a role for which *FOXO1* gene would be a relevant candidate [25, 14].
7. Finally, *G*9 is activated by *G*8 and inhibits death, a role known to be played by the antiapoptotic gene *Bcl-2* [20].

### 3.3 Population trajectory and differentiation states of a reference case

We first focus on the results obtained for a reference case, i.e. an initial population composed with 980 CD8 T cells and 700 APCs. All the parameters used to perform simulations can be found in Section 2.8. Figure 2 shows the evolution of the CD8 T lymphocyte population and of each sub-population of differentiated cells (Naive, Bipotent, Effector and Memory populations). Results represent average and standard deviations over 10 simulations (using different random generator seeds). The results capture the expected dynamics of the CD8 T cell population qualitatively, i.e. naive cells rapidly disappear, differentiating into bipotent cells that further generate effector or memory cells. In the effector state, cells proliferate rapidly, reaching a peak followed by a contraction phase, at which point memory cells appear and accumulate. In the last phase, only memory cells remain that form a stable population (see [54]).

**Figure 2:**
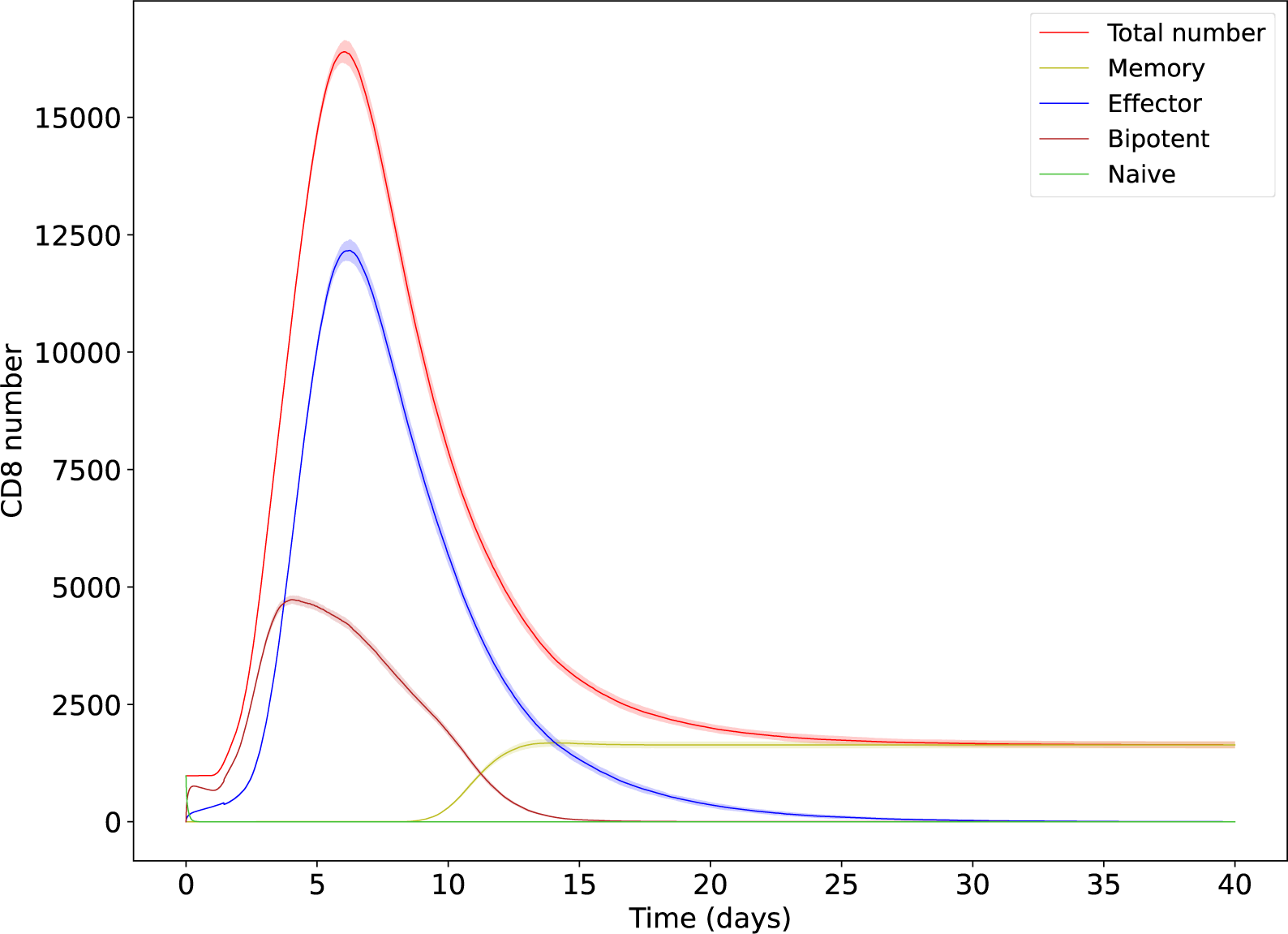
Time-dependent evolution of differentiation states of the CD8 T cell population over 10 simulations (mean (solid line) *±* std (shaded area)). The green, brown, blue, and yellow lines present the total number of naive, bipotent, effector, and memory phenotype of CD8 T cells, respectively, and the red line the total number of CD8 T cells.

From a more quantitative perspective, after 18 hours, all 980 naive cells initially present (green solid line) have become activated by contacting with APCs, 72% of the activated cells became bipotent cells (brown) while 28% differentiated into effector cells (blue). Bipotent cells gradually transform into effector cells, which proliferate very rapidly and reach a peak between days 5 and 10 post-immunization. On average, a total of 16, 422 T cells (red) were formed at the peak (a 16.8 fold expansion) and 1, 637 memory cells (yellow) remain in the population at the end of the simulation (40 days). For more details about cell numbers and standard deviations, see Table 6.

**Table 6:** Comparative values of different initial data, where protein degradation rate of *G*3 equals 0.021. All ratios are computed using mean values of specified quantities.

| T cell | APC | Ratio T cell/ APC | T cell at the peak (mean ± std) | Ratio T cell at the peak/ initial T cell | Time of the peak (days) (mean ± std) | Memory cell at the end (mean ± std) | Ratio Memory/ T cell at the peak (%) |
| --- | --- | --- | --- | --- | --- | --- | --- |
| 35 | 25 | 1.4 | 500 ± 37 | 14.3 | 7.2 ± 0.5 | 61 ± 11 | 12 |
| 70 | 50 | 1.4 | 1,135 ± 29 | 16.2 | 6.7 ± 0.2 | 121 ± 10 | 11 |
| 140 | 100 | 1.4 | 2,292 ± 48 | 16.4 | 6.4 ± 0.2 | 228 ± 26 | 10 |
| 210 | 150 | 1.4 | 3,469 ± 62 | 16.5 | 6.2 ± 0.1 | 357 ± 37 | 10 |
| 980 | 700 | 1.4 | 16,422 ± 235 | 16.8 | 6.0 ± 0.1 | 1,637 ± 72 | 10 |
| 1,960 | 1,400 | 1.4 | 32,802 ± 165 | 16.7 | 6.0 ± 0.1 | 3,150 ± 95 | 10 |
| 210 | 25 | 8.4 | 2,177 ± 128 | 10.4 | 7.4 ± 0.2 | 257 ± 19 | 12 |
| 210 | 50 | 4.2 | 2,924 ± 101 | 13.9 | 6.8 ± 0.2 | 286 ± 32 | 10 |
| 210 | 210 | 1.0 | 3,583 ± 45 | 17.1 | 6.2 ± 0.1 | 351 ± 44 | 10 |
| 210 | 420 | 0.5 | 3,596 ± 78 | 17.1 | 6.0 ± 0.1 | 395 ± 42 | 11 |
| 210 | 630 | 0.4 | 3,577 ± 57 | 17.0 | 6.0 ± 0.1 | 368 ± 24 | 10 |
| 980 | 117 | 8.4 | 12,724 ± 472 | 13.0 | 7.1 ± 0.1 | 1,337 ± 87 | 11 |
| 980 | 175 | 5.6 | 14,592 ± 426 | 14.9 | 6.7 ± 0.1 | 1,472 ± 79 | 10 |
| 980 | 1,050 | 0.9 | 16,502 ± 274 | 16.8 | 6.0 ± 0.1 | 1,686 ± 106 | 10 |
| 980 | 1,960 | 0.5 | 16,750 ± 229 | 17.1 | 6.0 ± 0.1 | 1,738 ± 76 | 10 |
| 980 | 2,800 | 0.4 | 16,744 ± 284 | 17.1 | 6.0 ± 0.1 | 1,733 ± 115 | 10 |

Noticeably, despite the stochasticity introduced at the molecular level with the mRNA bursting regime and at the cellular level with the random cell movement, the resulting output from 10 independent simulations is remarkably predictable, with little variation between two simulations of the model, characterized by small standard deviations (see Figure 2, where standard deviations are illustrated by narrow shaded areas around the mean, and Table 6).

### 3.4 Time-dependent evolution of mRNA and protein

Having demonstrated a robust behavior of the model at the cellular level, we then assess its behavior at the molecular level. For this we plot in Figure 3 the histograms over time of mRNA expression for the 9 core genes of the augmented GRN through one simulation randomly selected from the 10 simulations (Figure 2). The times at which those distributions are shown were chosen to capture the early phase of expansion, the contraction phase and were more widely spaced for the final memory generation phase. Altogether the genes display the expected dynamics. *G*1 shows a very brief period of activation, due to its early activation by APC signalling which stops after the first division. *G*2 shows a much more sustained period of activation during the expansion phase. *G*3 is mostly expressed during the contraction phase and both *G*2 and *G*3 expressions go towards 0 at the end of the simulation. The frequency of *G*4 mRNA expression follows the distribution of *G*2, because *G*2 activates *G*4. The expression of *G*7 and *G*8 mirrors that of *G*5 and *G*6, which are mutually exclusive as expected from their toggle switch connection.

**Figure 3:**
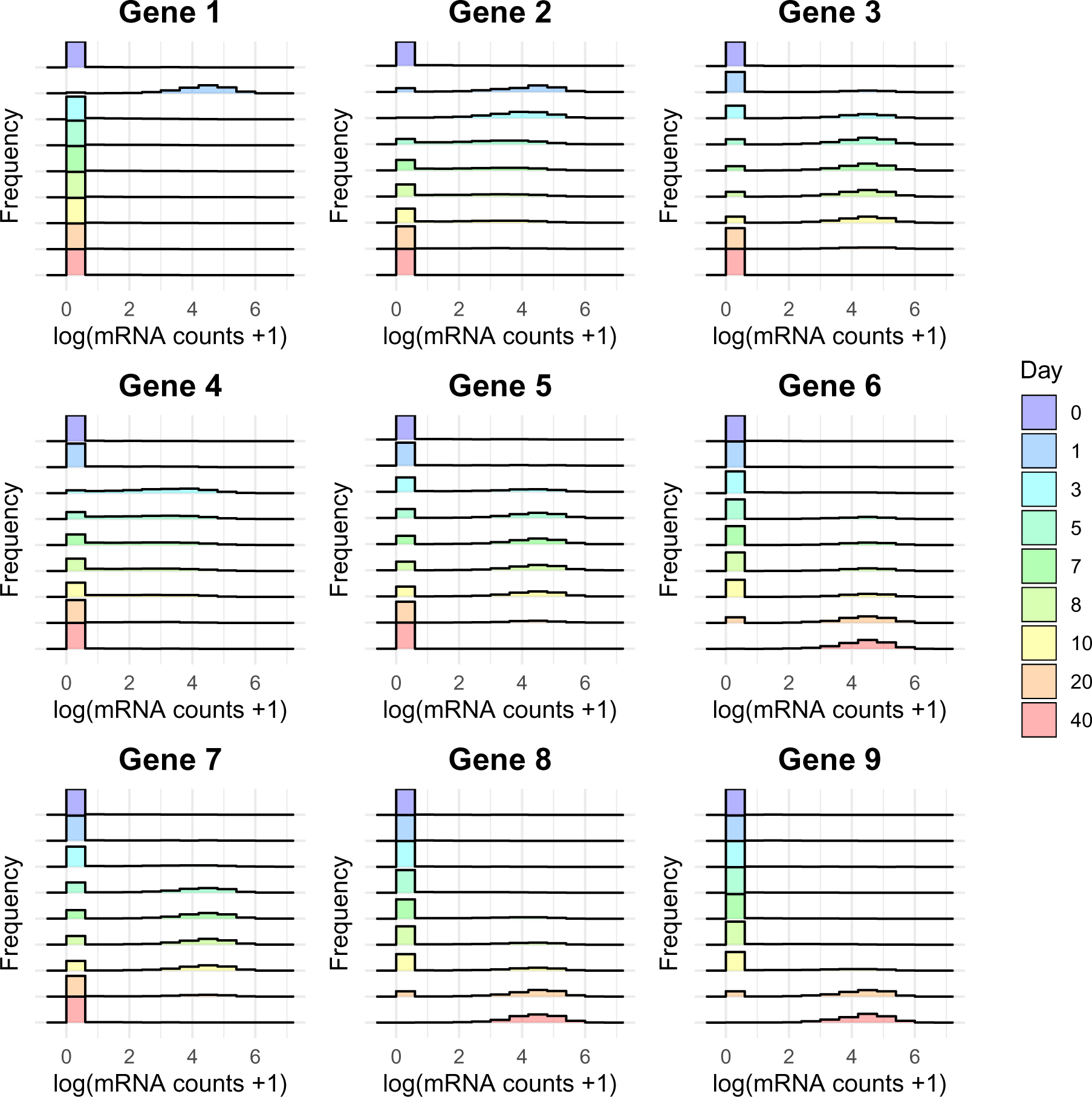
Time dependent evolution of mRNA distributions of 9 genes (see Figure 1A): x-axis shows the value of mRNA+1 in logarithmic scale, while y-axis shows the frequency observed at different time points.

It is important to note that those distributions harbour the characteristics known for patterns of gene expression at the single cell level, i.e. a strong zero component and a long tailed distribution. We have previously shown that the ability to generate realistic scRNA-seq datasets was a key asset of the bursty model [21].

We then explore the behaviour of the molecular model at the protein level. Figure 4 shows the time-dependent evolution of the mean expression of each protein across all cells, for the same simulation. Here too, the expected behaviour is observed.

**Figure 4:**
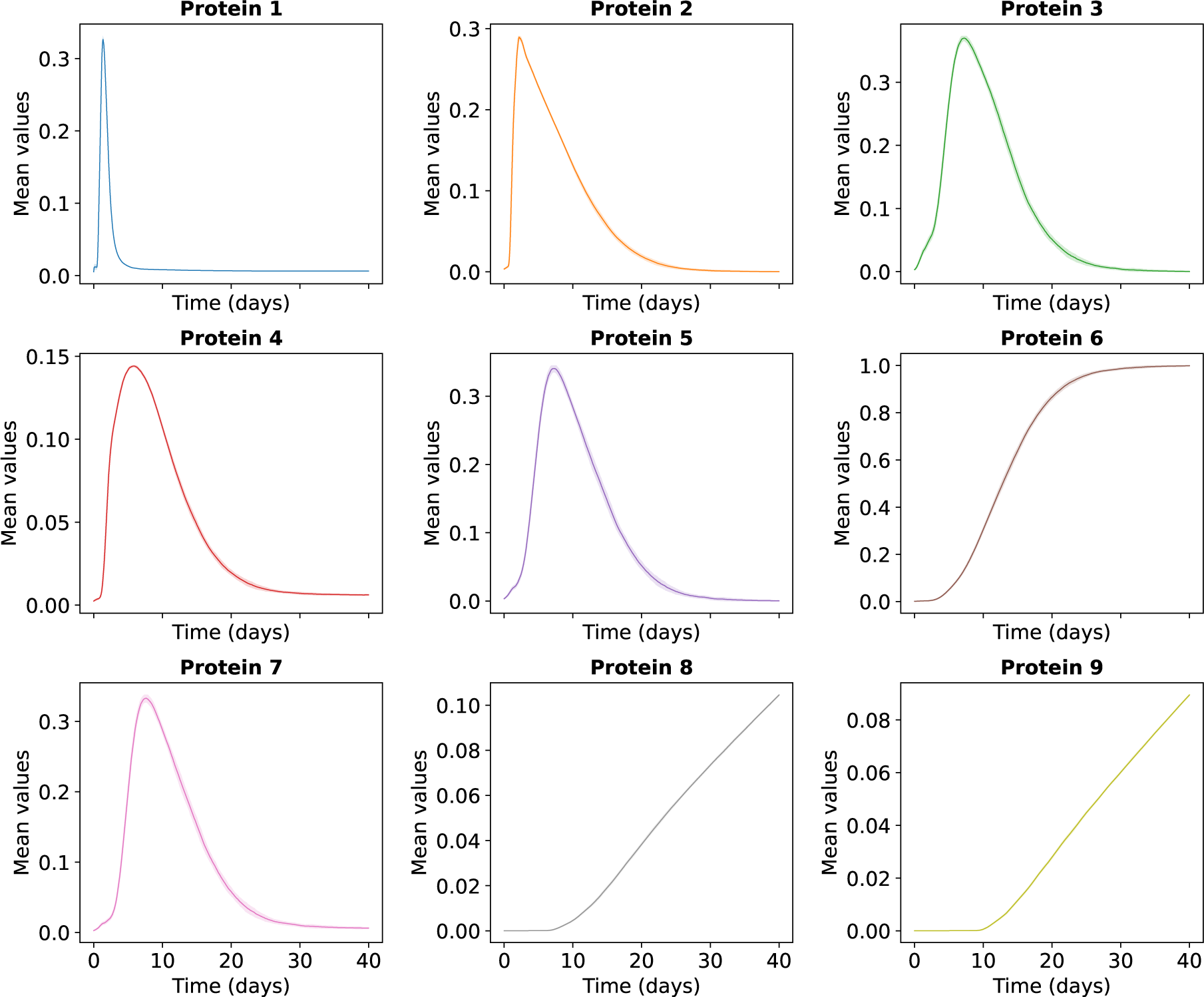
Time-dependent mean protein expressions over 10 simulations across all cells. Shown is the mean (solid line) *±* std (shaded area) for the 9 main genes of the augmented GRN.

The mean protein expression of *G*1 (blue line) first increases sharply, then decreases after 1 or 2 days, correctly representing the expected action of *G*1 in the GRN. While *P* 2 increases and quickly reaches a peak (orange line), *P* 3 slowly increases (green line) due to the inhibition of *G*3 by *G*2. When *P* 2 decreases then *P* 3 increases sharply and reaches a peak around the time of the response peak. Since *G*4 is activated by *G*2, the growth pattern of *P* 4 is similar to that of *P* 2. The toggle switch between *G*5 and *G*6 is visible in the mean protein expression between *P* 5 (purple) and *P* 6 (brown). This is mirrored in changes between *P* 7 and *P* 8, with the expression of *G*7 (pink line) increasing initially and decreasing as *G*8 increases. Similarly, *G*8 activates *G*9, then the yellow line of *G*9 rises following the increase of *G*8.

One should also notice that the time-dependent evolution of the average protein amount is associated with small standard deviations (represented by very narrow shaded areas around the mean), hence highlighting a pretty reproducible behaviour of the model.

### 3.5 UMAP analysis

The highly dimensional nature of scRNAseq data has called for the development of suitable dimensionality reduction techniques. Among those, the UMAP representation (Uniform Manifold Approximation and Projection [6]) has established itself as one of the most popular. We therefore assess the ability of the model to produce a relevant UMAP representation of the CD8 differentiation sequence (Figure 5). Our initial attempt at obtaining a UMAP representation of the model output based on the principle-based GRN gave rise to not very realistic images (see Supplementary Figure 4). At that stage we reasoned that the amount of information provided to the UMAP algorithm might have been too sparse. We therefore decided to add “decorating genes” (see Section 3.1) and used the augmented GRN instead. The decorating genes do not participate in the dynamics of the network, but add some redundant information that proved to be required to obtain the final correct UMAP representation observed on Figure 5.

**Figure 5:**
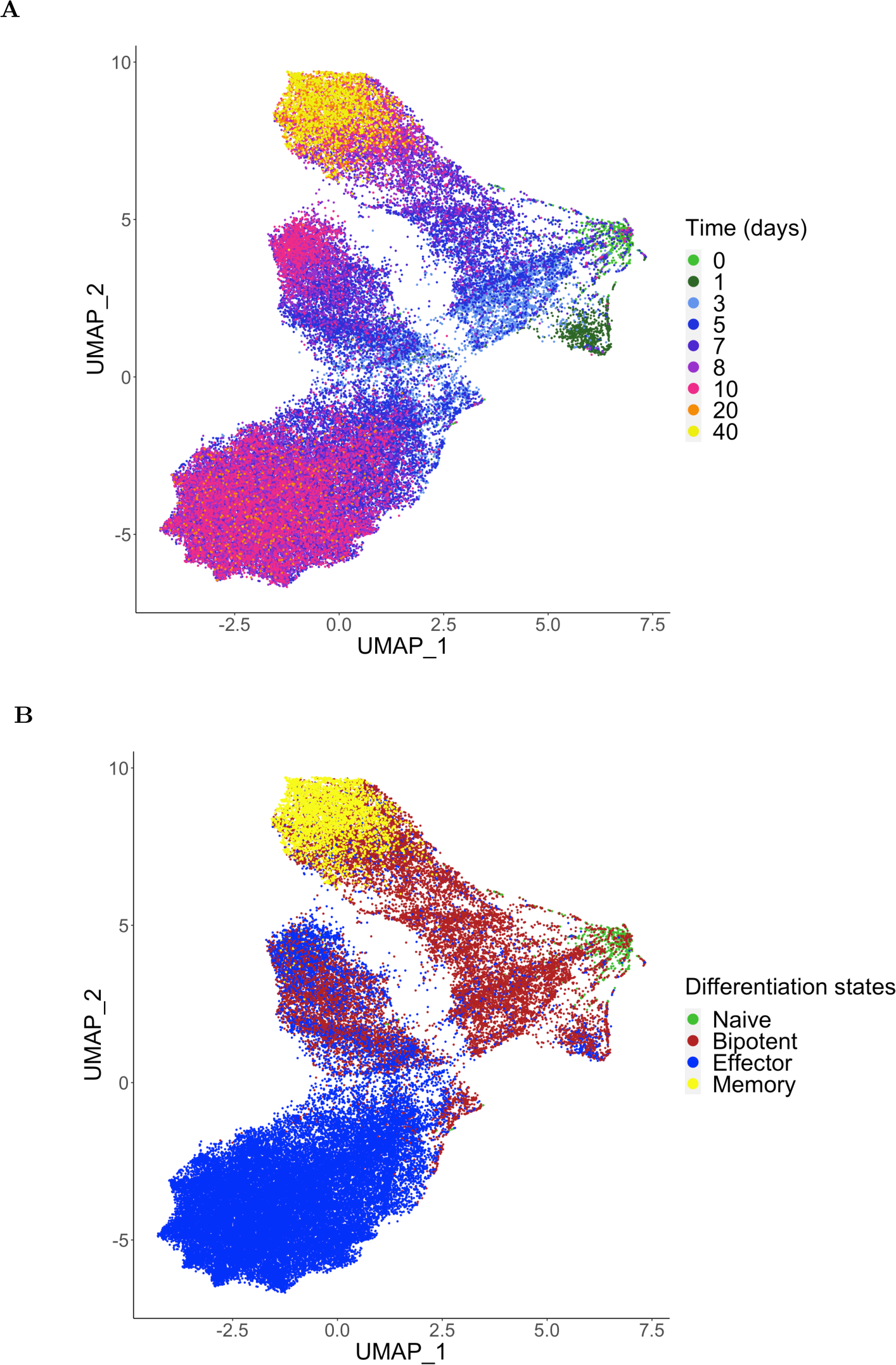
UMAP representation of CD8 T cell population dynamics over time. (A) Cells are color-coded as a function of the time they were observed. The same time points are displayed as in Figure 3. (B) The same graph, where the cells are now color-coded according to their phenotype.

On Figure 5A, one can clearly see the time-dependent trajectory evolution of the CD8 T cell population, starting at day 0 on the right and going to day 40 to the top left. The UMAP representation therefore reveals a correct temporal arrangement of the cells.

Figure 5B presents the trajectory evolution of the CD8 T cell population as a function of their differentiation states. We can see that at day 0 most cells are naive cells (green in both figures) clustered close together, since their initial molecule content is almost 0. Bipotent cells appear at an early stage (brown in 5B)) and then become effector or memory cells, giving rise to two branches of cells. This clearly shows the correct behavior of the *G*5/*G*6 toggle switch pushing cells out of their bipotent state and forcing them into an effector or memory fate. The vast majority of cells are effector cells (blue in 5B) around day 6, but there is no effector cell left by day 40, suggesting that they have gone through the expected contraction phase. Indeed, at the end of the simulation, i.e. on day 40 (yellow color code in 5A), only memory cells remain.

### 3.6 Sensitivity to parameter values variations

To analyze the impact of some parameters on the model behavior, we first assess the role played by the initial number of cells. Table 6 shows the effect of varying simultaneously the initial T cell number and the initial APC number, while keeping the same initial T cell/APC ratio, or keeping the initial T cell number constant (either 210 or 980) and varying the initial APC number. We measure the impact of varying those numbers on five variables:

1. The number of T cells at the peak of the response;
2. The ratio between the average number of T cells at the peak of the response and the initial number of T cells;
3. The time of the response peak;
4. The average number of memory T cells at the end of the simulation;
5. The ratio between the average number of memory T cells at the end of the simulation and the average number of T cells at the peak of the response.

We first explore the impact of modifying initial cell counts while keeping a constant ratio of T cells to APCs (equal to 1.4) on the overall amount of cells. One can see in Table 6 that there is a steady monotonous increase in the number of cells at the peak. Nevertheless if one examines the ratio between the average number of T cells at the peak of the response and the initial number of T cells, the situation is very different as illustrated in Figure 6A. There is initially an increase in this variable, which then stabilizes in a very narrow range between 16.2 and 16.8. This amplification ratio therefore appears as a relatively robust emerging property of the model and suggests a minimal initial cell density is required for the optimal expansion of the effector population, as observed *ex vivo* [43].

**Figure 6:**
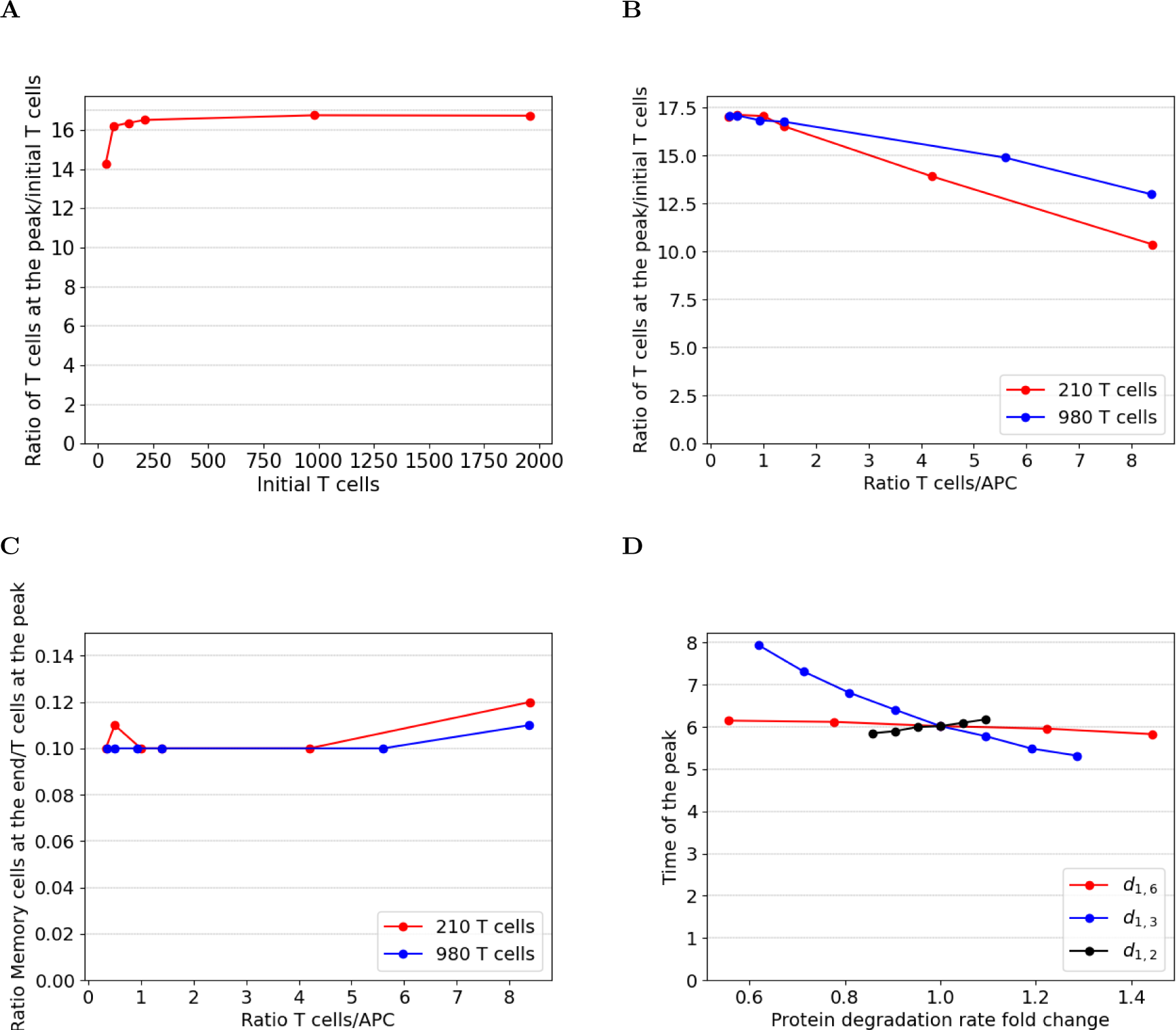
Comparison of variables of interest when varying initial cell counts. (A) Average cell expansion at the peak of the response as a function of initial cell numbers over 10 simulations. (B) Average cell expansion at the peak of the response as a function of the ratio of initial T cells to APCs counts, in two cases comprising 210 and 980 T cells. (C) Ratio of the average memory cell counts at the end of the simulation to average T cells count at the peak, as a function of the ratio of initial T cells to APCs counts, in two cases comprising 210 and 980 T cells. (D) Effect of proteins 2, 3 and 6 degradation rates upon the time of the peak (for a Ratio T cells/APC equals 1.4). Reference values of protein degradation rates are given in Table 3.

One should note that the maximum number of cells generated was more than 32, 000 cells, high- lighting the ability of the model to generate very large amounts of cells. The computational time was only 2h in this case (see Section 2.5).

We then explore the impact of varying the initial number of APCs while keeping constant the initial number of T cells (210 or 980). Larger ratios yield weaker cellular expansions in both situations (Figure 6B), highlighting a positive effect of the antigen load on CD8 T cell expansion [38]. The range of values we used indicates there may be an optimal ratio of 2 APCs for 1 T cell, when 980 CD8 T cells are initially seeded.

We finally can demonstrate that the ratio between the average number of memory T cells at the end of the simulation and the average number of T cells at the peak of the response is quite insensitive to the initial APCs to T cells ratio (Figure 6C), highlighting the initial antigen load does not influence the extent of cellular contraction after the peak of the response, as demonstrated *in vivo* [4]. It turns out that the value of this variable proved to be extremely robust, around 10% *−* 12% in all of the situations tested (see Table 6), which corresponds to *in vivo* situations [39].

Investigating the impact of molecular processes on the dynamics of the immune response through variations of different parameters (see Section 2.10) showed, as expected, that variations of Protein 2 degradation rate mostly impact the production of T cells at the peak of the response while variations of Protein 6 degradation rate mostly impact the generation of memory cells (Supplementary Figure 5). However, although the time at which the peak is observed is around 6-7 days and does not seem sensitive to initial cell counts, we could demonstrate the role played by the degradation rate of Protein 3 in setting the time of the response peak (Figure 6D and Table 7). Indeed the time at which the expansion phase reaches its peak value is monotonically decreasing as a function of the degradation rate.

**Table 7:** Effects of protein degradation rates *d*_1_*_,·_* of genes 2, 3 and 6 on various variables of interest. In each case, initial cell counts are: 980 T cells and 700 APC (ratio equals 1.4).

| Value | T cell at the peak (mean ± std) | Ratio T cell at the peak/ initial T cell | Time of the peak (days) (mean ± std) | Memory cell at the end (mean ± std) | Ratio Memory/ T cell at the peak (%) |
| --- | --- | --- | --- | --- | --- |
| $d_{1,2}$ | | | | | |
| 0.023 | 19,821± 190 | 20 | 6.2± 0.1 | 1,918± 82 | 10 |
| 0.022 | 18,029± 150 | 18.4 | 6.1± 0.1 | 1,783± 68 | 10 |
| 0.021 | 16,422 ± 235 | 16.8 | 6.0 ± 0.1 | 1,637 ± 72 | 10 |
| 0.02 | 14,944± 150 | 15.3 | 6.0± 0.1 | 1,493± 93 | 10 |
| 0.019 | 13,453± 138 | 13.7 | 5.9± 0.1 | 1,371± 64 | 10 |
| 0.018 | 12,125± 101 | 12.4 | 5.9± 0.1 | 1,205± 51 | 10 |
| $d_{1,3}$ | | | | | |
| 0.027 | 12,941 ± 120 | 13.2 | 5.3±0.1 | 1,582 ± 46 | 12 |
| 0.025 | 13,899 ± 110 | 14.2 | 5.5± 0.1 | 1,601± 57 | 12 |
| 0.023 | 15,105 ± 173 | 15.4 | 5.8 ± 0.1 | 1,632 ± 47 | 11 |
| 0.021 | 16,422 ± 235 | 16.8 | 6.0 ± 0.1 | 1,637 ± 72 | 10 |
| 0.019 | 17,400 ± 684 | 17.7 | 6.4 ± 0.1 | 1,592 ± 76 | 9 |
| 0.017 | 19,107 ± 600 | 19.5 | 6.8 ± 0.1 | 1,562 ± 137 | 8 |
| 0.015 | 21,305 ± 466 | 21.7 | 7.3 ± 0.1 | 1,672 ± 84 | 8 |
| 0.013 | 23,857 ± 258 | 24.3 | 7.9 ± 0.1 | 1,746 ± 87 | 7 |
| $d_{1,6}$ | | | | | |
| 0.013 | 15,433± 162 | 15.8 | 5.8± 0.1 | 1,983± 98 | 13 |
| 0.011 | 15,990± 88 | 16.3 | 6.0±0.1 | 1,857± 40 | 12 |
| 0.009 | 16,422 ± 235 | 16.8 | 6.0 ± 0.1 | 1,637 ± 72 | 10 |
| 0.07 | 16,620± 155 | 17.0 | 6.1 ±0.1 | 1,312± 81 | 8 |
| 0.05 | 16,881± 109 | 17.2 | 6.2± 0.1 | 785± 43 | 5 |

We can therefore show the functional impact of a molecular parameter value of the molecular GRN at the cell population level: a higher degradation rate for the protein controlling cell death induces an earlier peak in the response. The accumulation of *P* 3 up to its threshold value to induce death takes longer time with a more unstable protein, suggesting a higher degradation rate allows the expanding population to reach its maximum more rapidly (even though this earlier peak is associated with less T cells at the peak, see Supplementary Figure 5 and Table 7), although more complex explanations might be at stake. This is however a very interesting result which illustrates the benefit of this model, since it can only be obtained with the proper coupling of all scales and the inclusion of a mechanistic gene model.

## 4 Discussion

In this work, we have presented a multiscale model of the CD8 T cell immune response, from naive to memory cells, in which cell behavior (proliferation, apoptosis or differentiation) is determined by molecular intracellular dynamics. We have demonstrated that this modeling approach can be used to correctly reproduce the expected population dynamics of CD8 T cells [54], using a 9-component principle-based GRN.

The use of a mechanistic model of gene expression, the two-state model in its bursty regime [22], allowed us to obtain at all time points a realistic distribution of the amount of mRNAs. It is important to point out that these types of expression distributions are the main output of scRNAseq experiment and therefore comparing distributions represent a natural way to compare a model output to experimentally observed values [58]. The model-generated distributions can be used for comparison with experimental distributions, for instance using the Kantorovitch distance [9]. Interestingly, the output of the model makes it also possible to extract single-cell proteomics, therefore while it is currently challenging to perform single-cell proteomics [31] we can make predictions using our modeling and simulation framework.

UMAP representation is also a natural way of comparing the output of a model and an experimental single-cell dataset (see e.g. [58]). In our case, the GRN we introduced was able to correctly generate a time-dependent evolution of single cells at the molecular level, and we could observe the expected time-dependent trajectory in the UMAP space, as well as the branching resulting from the decision making process through the toggle switch as observed experimentally [54].

One of the main difficulties we encountered deals with the parameterization of the model. Numerous parameters have to be defined both at the molecular and at the cellular levels (see Section 2). Since the aim of this work was to establish a proof-of-concept that we could reproduce expected dynamics, parameters were tuned for such a purpose. In trying to improve the realism of our model, more values should be estimated from experimentally measured datasets. However, it is a notoriously difficult task to obtain realistic values for certain molecular parameters, such as degradation rates, from experiments [47].

Although a full sensitivity analysis was beyond the scope of the present work, we nevertheless could analyze the impact of a few parameters. Quite interestingly, some characteristic population-level behavior of the model proved to be very robust to specific influences. Besides robust parameters, there are also some for which a sensitivity analysis will be required, such as the interaction strength of the toggle switch between *G*5 and *G*6, or the time-dependent parameters for cell-cell interaction. Preliminary testing shows that varying those parameters may well change the population dynamics, as well as the differentiation states.

Such a sensitivity analysis will be facilitated by two important characteristics of the model:

1. The extremely low variation between two repeats will be critical, allowing us to reduce the amount of necessary repeats to run them in parallel on a computing cluster.
2. Concerning the runtime in Simuscale , it is significantly faster and more competitive than previous software. The most extensive task (simulating 32,000 cells at the peak of the response) took 62min to be executed on an Apple M2 Pro with 32GB of memory.

Once the parameters had been calibrated, the model proved to be able to reconstruct the temporal dynamics of the CD8 T cell population in general, and of each differentiation state in particular.

The next step in our approach will consist in changing the principle-based GRN to a more realistic one. We are currently in the process of using CARDAMOM [58] to infer a GRN from the time-stamped

scRNAseq data from [34]. Interestingly, the model described in the present work already gives us an idea of 1) what to expect, i.e. we need at least a toggle switch, which will lead to the differentiation state of the cell; and 2) where the signals should affect GRN behavior, i.e. APC signalling should influence the proliferation, and TCC signalling should impact the death. Moreover, our experience in choosing the degradation rate parameter values will be useful too, as long as they have not been experimentally determined. Finally in the case where we will try to fit experimental data, the mRNAs distributions observed experimentally will be compared directly to the model output, and the model parameters will be adjusted to minimize any discrepancy.

There are a number of changes that can be envisioned to improve the modeling approach:

1. The 3D domain represented in Simuscale is currently a fixed-size cube, which can be reduced to speed up the contraction phase. It could also be enlarged or its shape be modified, to assess the influence on the population dynamics.
2. All cellular decisions are built upon the crossing of a given threshold level. We could assess the impact of changing this rule for a more realistic one that would make the *probability* of a decision to depend upon a given protein level.
3. As of now, signalling can only be modeled by cell-to-cell contact. However, we have previously demonstrated the impact of the local diffusion of IL2 on the CD8 T cell response [18]. We could therefore implement a diffusion process in the extracellular space, although it would have to be carefully drafted not to slow down the code execution too significantly.

## Acknowledgments

T.N.T.N and M.M. have been funded through the ARN MEMOIRE project (MEMOIRE ANR-18- CE45-0001). We thank all members of the project for helpful discussions. We also thank Ulysse Herbach (Inria Nancy) who was very helpful to implement the bursty regime model within Simuscale and the P^ole Scientifique de Modélisation Numérique (PSMN) of ENS de Lyon for providing the computational resources. We also thank the BioSyL Federation and the LabEx Ecofect (ANR- 11-LABX-0048) of the University of Lyon for inspiring scientific events.

## Author Contributions

Conceptualization, T.N.T.N, C.A., O.G and F.C; Funding acquisition, O.G and F.C.; Investigation, All; Methodology, T.N.T.N, O.G and F.C; Project administration, O.G and F.C.; Software, T.N.T.N,

M.M., S.B.; Supervision, O.G and F.C.; Visualization, T.N.T.N, O.G and F.C; Writing – original draft, T.N.T.N, O.G, and F.C.; Writing – review & editing, All.

## Supplementary Figures

**Supplementary Figure 1:**
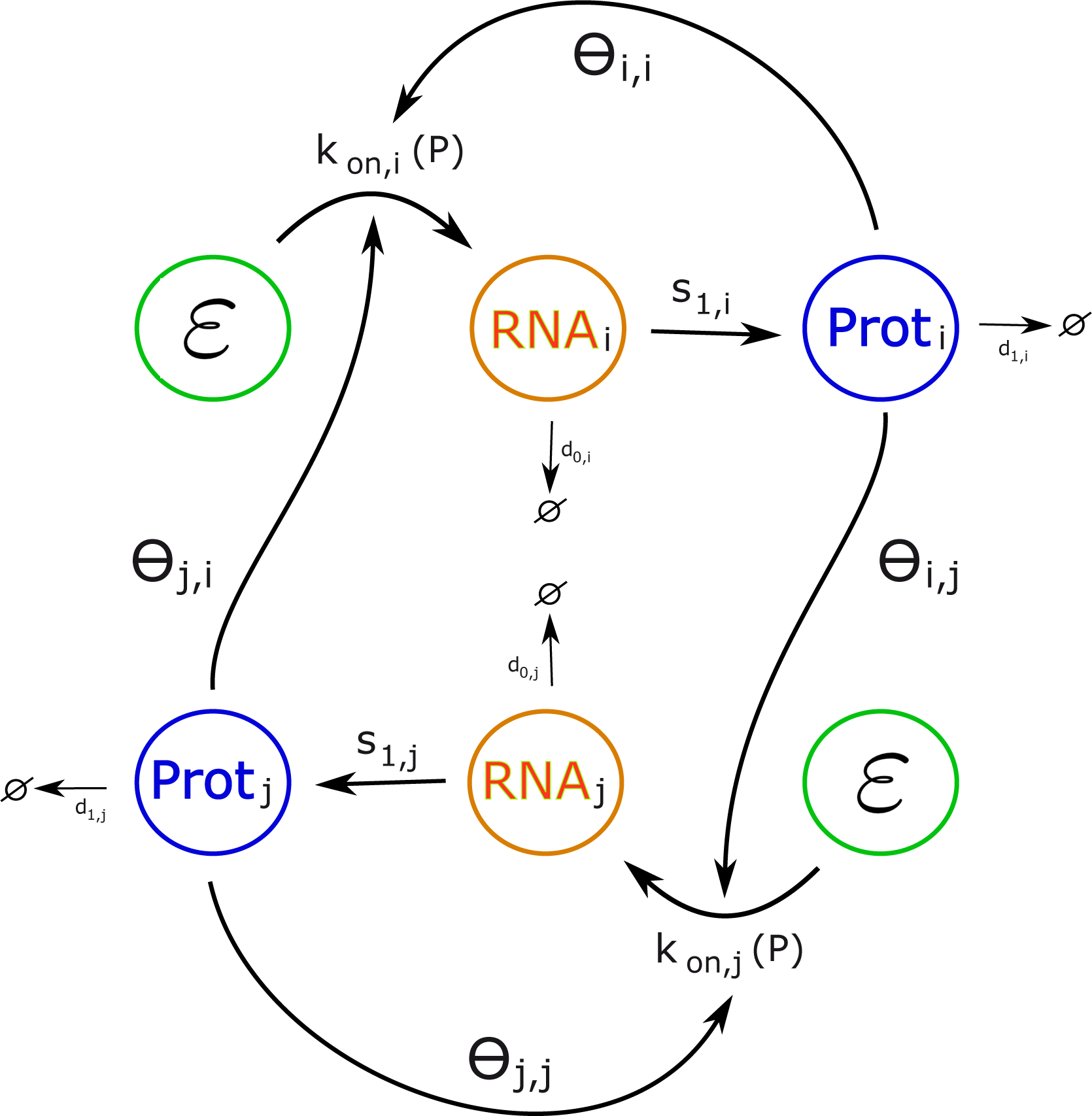
A schematic representation of a GRN connecting genes in the bursty regime. Two genes *G_i_* and *G_j_* are represented.

**Supplementary Figure 2:**
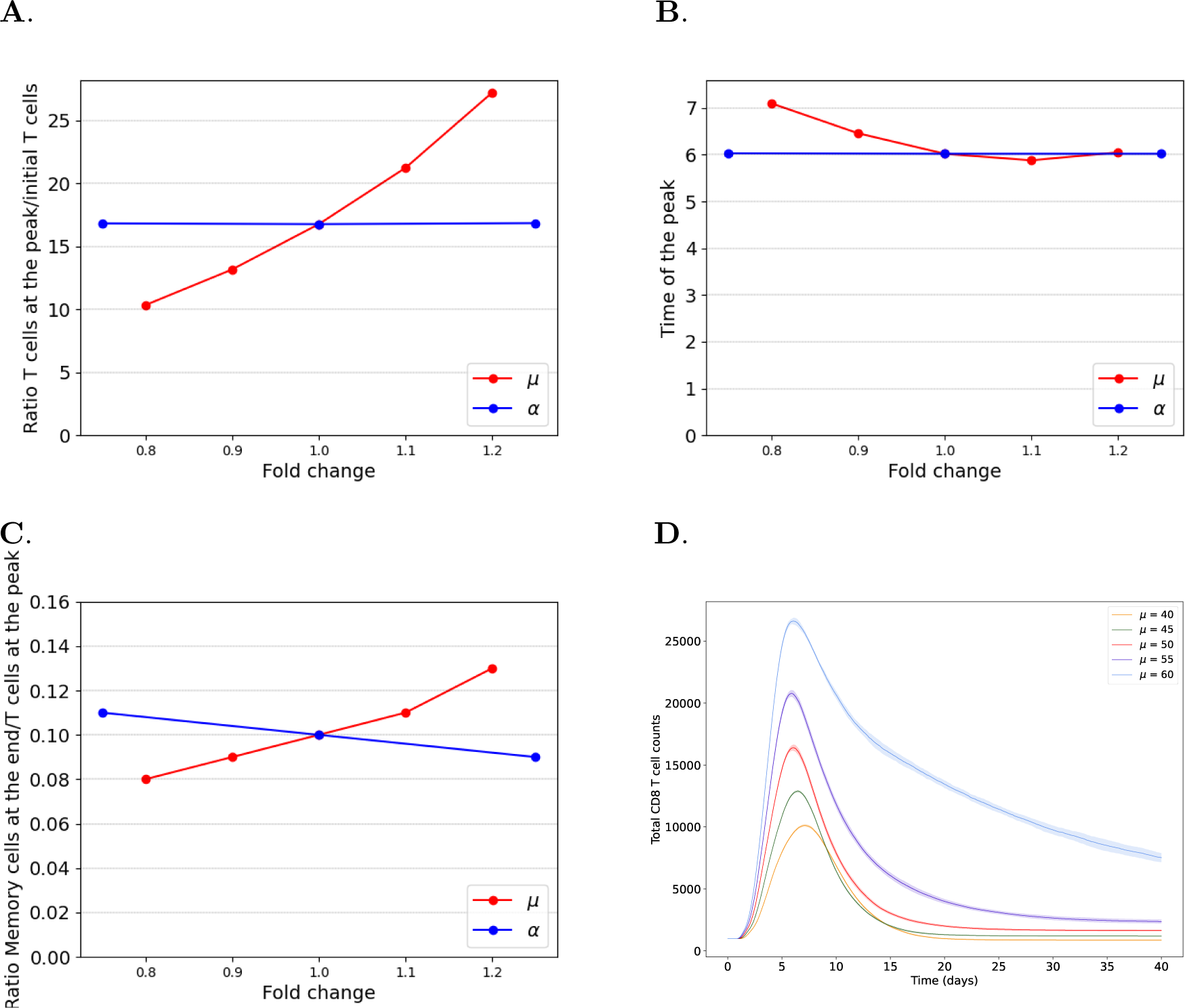
Effect of variations of parameters *α* and *µ* on variables of interest. (A) The ratio between CD8 T cells at the peak of the response and the initial T cell number. (B) The time of the peak. (C) The ratio between memory T cells at the end of the simulation and the number of T cells at the peak of the response. Reference values are *α* = 20 and *µ* = 50, as indicated in the text. (D) Evolution of the total population of CD8 T cells for several values of *µ*, corresponding to *±*20% of the reference value *µ* = 50.

**Supplementary Figure 3:**
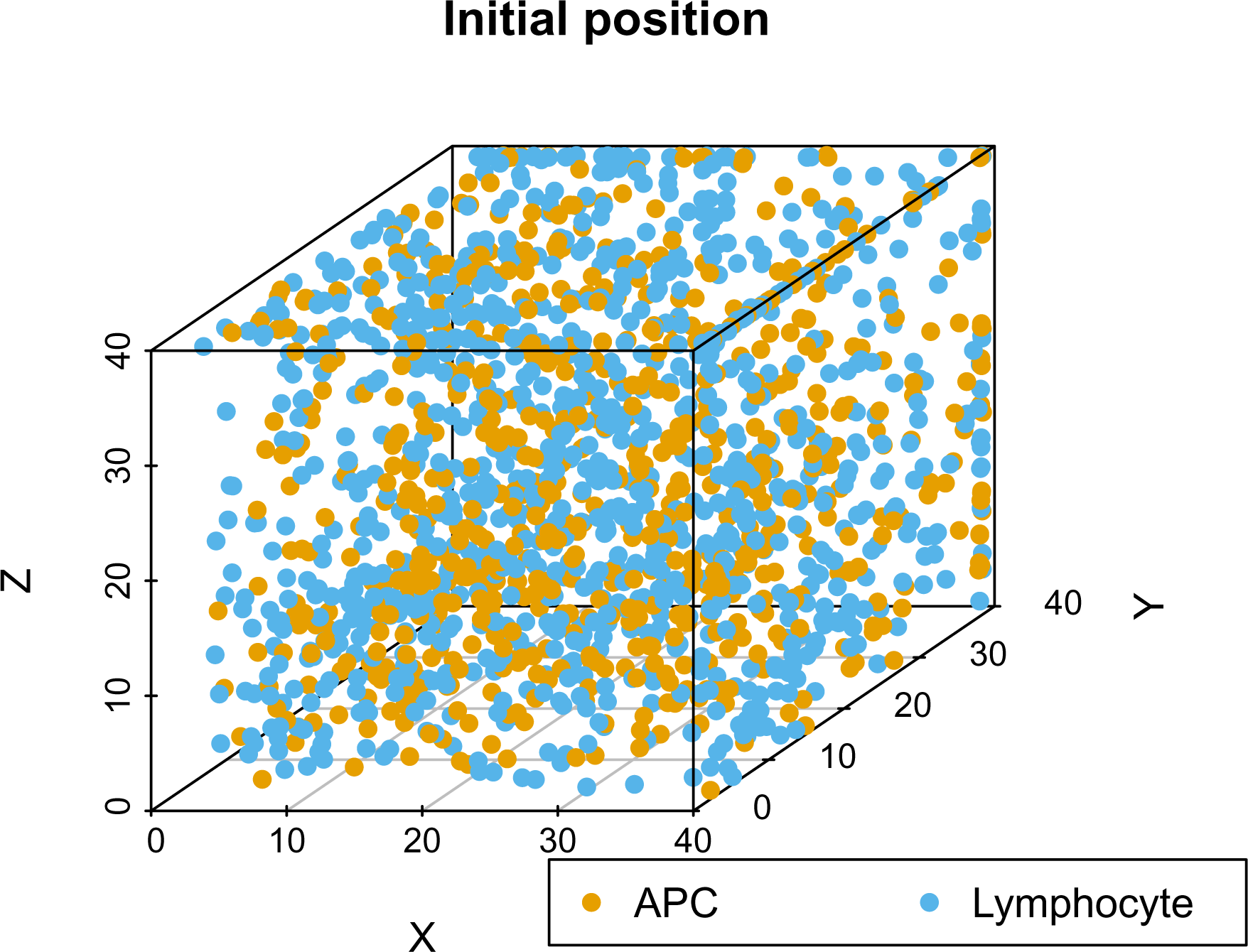
Random positions of cells at the beginning of a simulation.

**Supplementary Figure 4:**
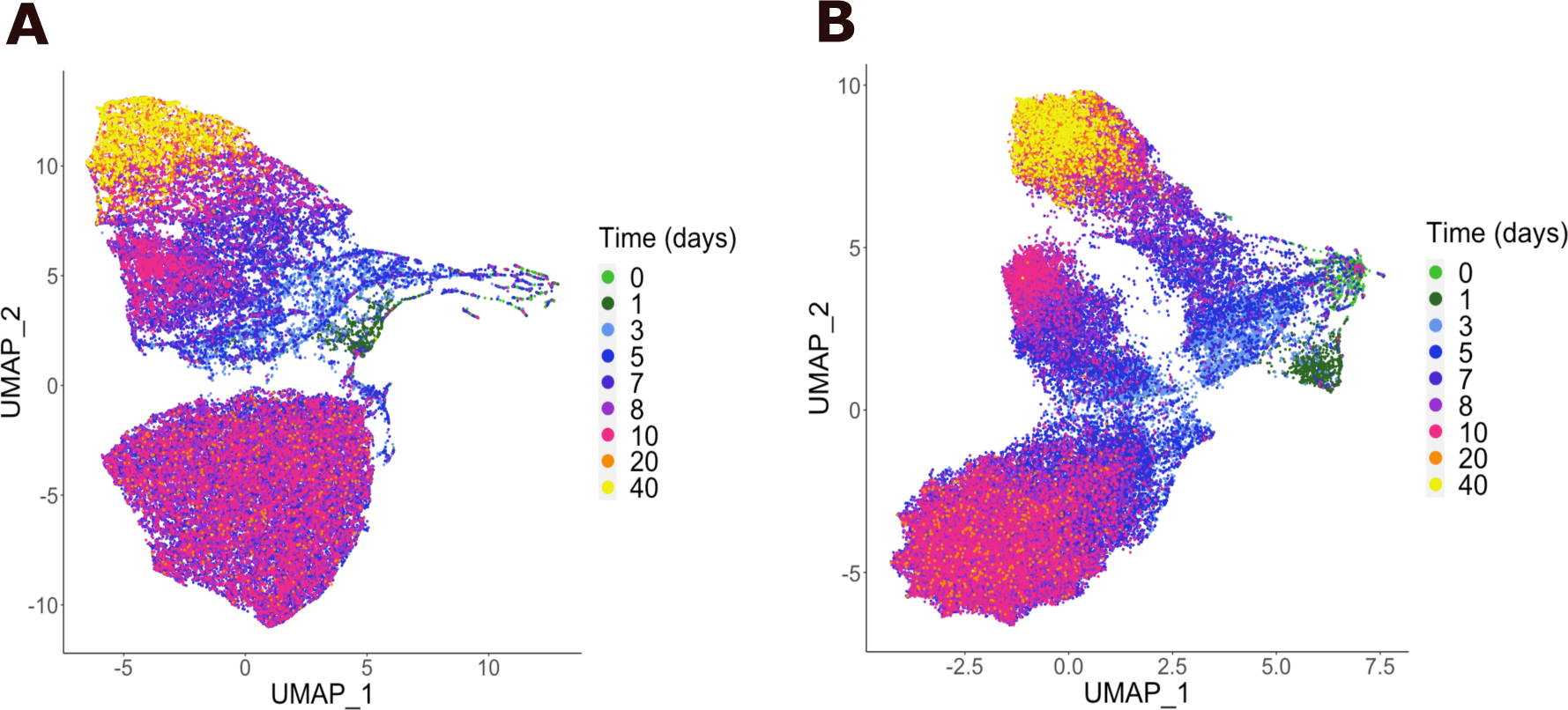
UMAP representation of CD8 T cell population dynamics over time using either (A) only the 9-genes principle-based GRN, or (B) the augmented GRN, based on the 9 core genes plus the 27 “decorating” genes. The figure in (B) is repeated here from Figure 5A, to allow direct comparison.

**Supplementary Figure 5:**
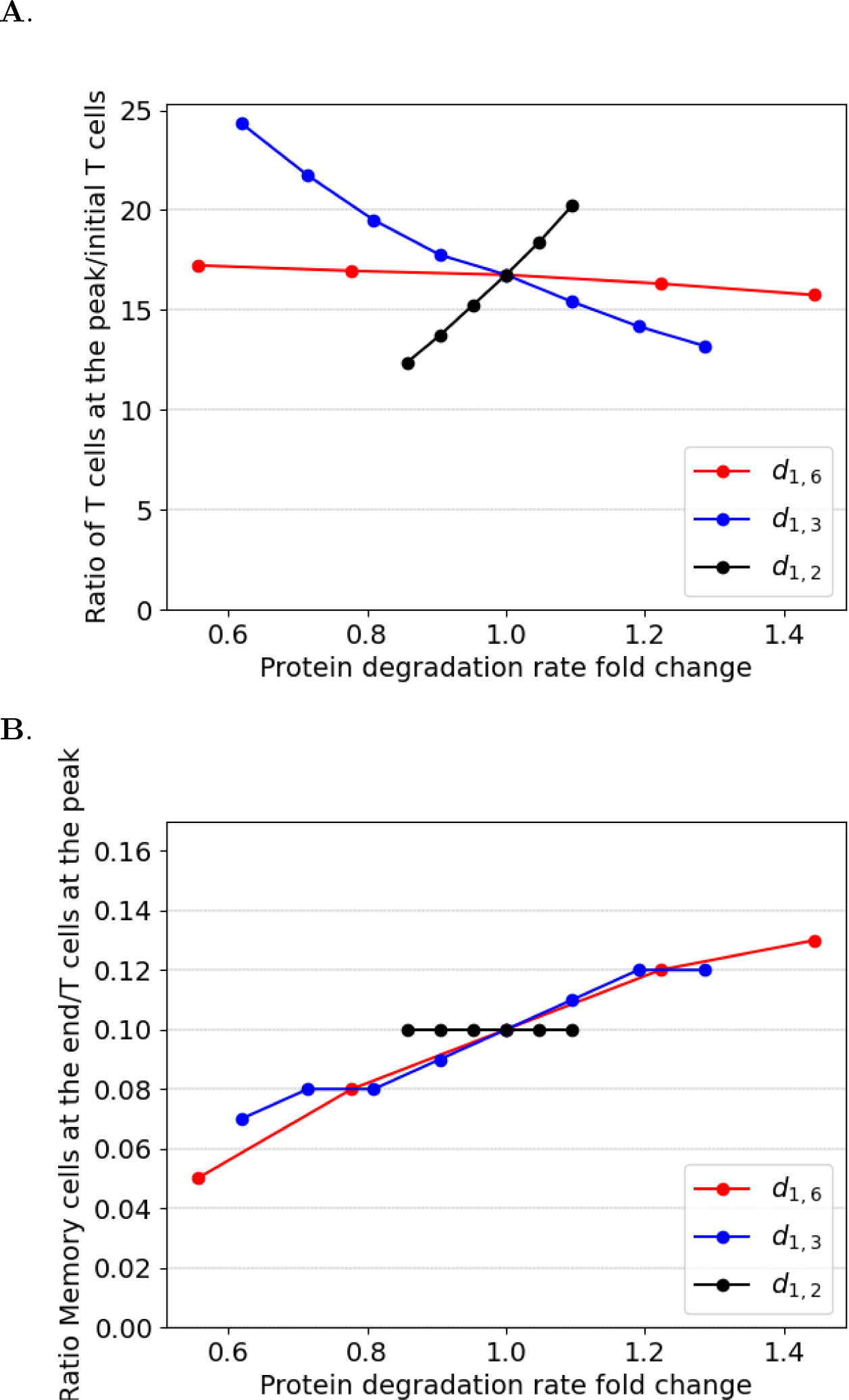
Effect of proteins 2, 3 and 6 degradation rates on variables of interest: (A) the ratio between CD8 T cells at the peak of the response and the initial T cell number, (B) the ratio between memory T cells at the end of the simulation and the number of T cells at the peak of the response. In both cases, the ratio T cells/APC equals 1.4, and reference values for protein degradation rates are given in Table 3.

